# Integrated Fluorescence Microscopy (iFLM) for Cryo-FIB-milling and In-situ Cryo-ET

**DOI:** 10.1101/2023.07.11.548578

**Authors:** Jae Yang, Veronika Vrbovská, Tilman Franke, Bryan Sibert, Matthew Larson, Alex Hall, Alex Rigort, John Mitchels, Elizabeth R. Wright

## Abstract

Correlative cryo-FLM-FIB milling is a powerful sample preparation technique for *in situ* cryo-ET. However, correlative workflows that incorporate precise targeting remain challenging. Here, we demonstrate the development and use of an integrated Fluorescence Light Microscope (iFLM) module within a cryo-FIB-SEM to enable a coordinate-based two-point 3D correlative workflow. The iFLM guided targeting of regions of interest coupled with an automated milling process of the cryo-FIB-SEM instrument allows for the efficient preparation of 9-12 ∼200 nm thick lamellae within 24 hours. Using regular and montage-cryo-ET data collection schemes, we acquired data from FIB-milled lamellae of HeLa cells to examine cellular ultrastructure. Overall, this workflow facilitates on-the-fly targeting and automated FIB-milling of cryo-preserved cells, bacteria, and possibly high pressure frozen tissue, to produce lamellae for downstream cryo-ET data collection.

## Introduction

Biological systems are complex, understanding the intricate interactions which enable life requires multiplexed analytical approaches. One structural biology technique that has risen to prominence for cell biology research is cryo-electron tomography (cryo-ET). Cryo-ET is used for the visualization of biological machinery *in situ* at the molecular level, giving us insights into the complex interactions occurring in individual cells (1). Due to electron beam penetration limitations of the TEM, in addition to cryo-ultramicrotomy (2), samples can be thinned via cryo-focused ion beam (cryo-FIB) milling (3) to prepare thin (< 300 nm) cryo-lamellae suitable for cryo-ET. While cryo-ET is powerful for visualizing the cogs of the biological machine, it remains challenging to determine the location of specific entities, especially when their structures are not known. Traditionally, fluorescent, multi-color labeling is used to identify macromolecules, complexes, and organelles, and this type of labeling also supports colocalization imaging experiments where two or more fluorophores can be visualized to reveal their respective positions in 3D. However, fluorescence light microscopy (FLM) does not provide high-resolution structural information. Correlative light and electron microscopy (CLEM) technologies allow us to combine the benefits of LM and cryo-ET to locate unknown structures quickly and efficiently, and then recall them in the TEM at high-resolution (4, 5). Thus, FLM-guided milling and correlative cryo-ET workflows facilitate the 3D mapping of both the labelled structure and its immediate sociology (6). Despite great efforts (6-10) to advance correlation and targeting approaches, correlative cryo-FLM-FIB suffers from low accuracy in z-axis positioning of regions of interest (ROI), and requires optimized fiducial markers and complicated 3D correlative algorithms, leading to low throughput. Here, we illustrate the requirements needed for successful 3D ROI relocation and demonstrate a practical and adaptable approach compatible with integrated correlative multi-modal microscopy methods.

We developed a reliable and more streamlined 3D correlative cryo-FLM-FIB-ET workflow that combines precise ROI targeting and automated cryo-FIB milling using a commercially available integrated fluorescence light microscopy (iFLM)-Aquilos 2 cryo-FIB-SEM system (Thermo Fisher Scientific, Fig. 1, SI Appendix Figs. S1 and S2), and the commercial Maps and AutoTEM cryo (Thermo Fisher Scientific) software packages, both of which are popular and accessible to the cryo-ET community. To avoid complicated transformation algorithms, we introduced a straightforward single-pair ROI-fiducial marker registration strategy (Fig. 2, SI Appendix Fig. S3 A-C). This approach applied a linear transformation to orient and register perpendicular SEM and FIB views acquired at different angles. This information was merged with the iFLM image stacks to update the z-location of the ROI relative to the fiducial marker. We also demonstrated the use of a sharp-cornered FIB-milled pattern (e.g., L-shaped pattern) as a fiducial marker imaged in reflection mode near the ROI (SI Appendix Fig. S4). We applied this workflow to identify and locate, in 3D, lipid droplet (LD)-mitochondria interactions in HeLa cells (Figs. 3 and 4), to routinely produce 200 nm thin lamella for cryo-ET analysis (Figs. 3 and 4, SI Appendix Figs. S5 and S6). This 3D multi-modal correlative workflow is compatible with correlation data from other FLM instruments and software solutions including CorRelator (5) that support cryo-ET data collection via CorRelator-SerialEM pipelines (11). Overall, our approach is an adaptable correlation technique for preparing ROI-containing cryo-lamellae from vitrified samples for *in situ* cryo-ET (12-15). We believe this workflow will improve applicability and throughput of cryo-FIB-milling techniques for structural cell biology.

**Figure 1.**
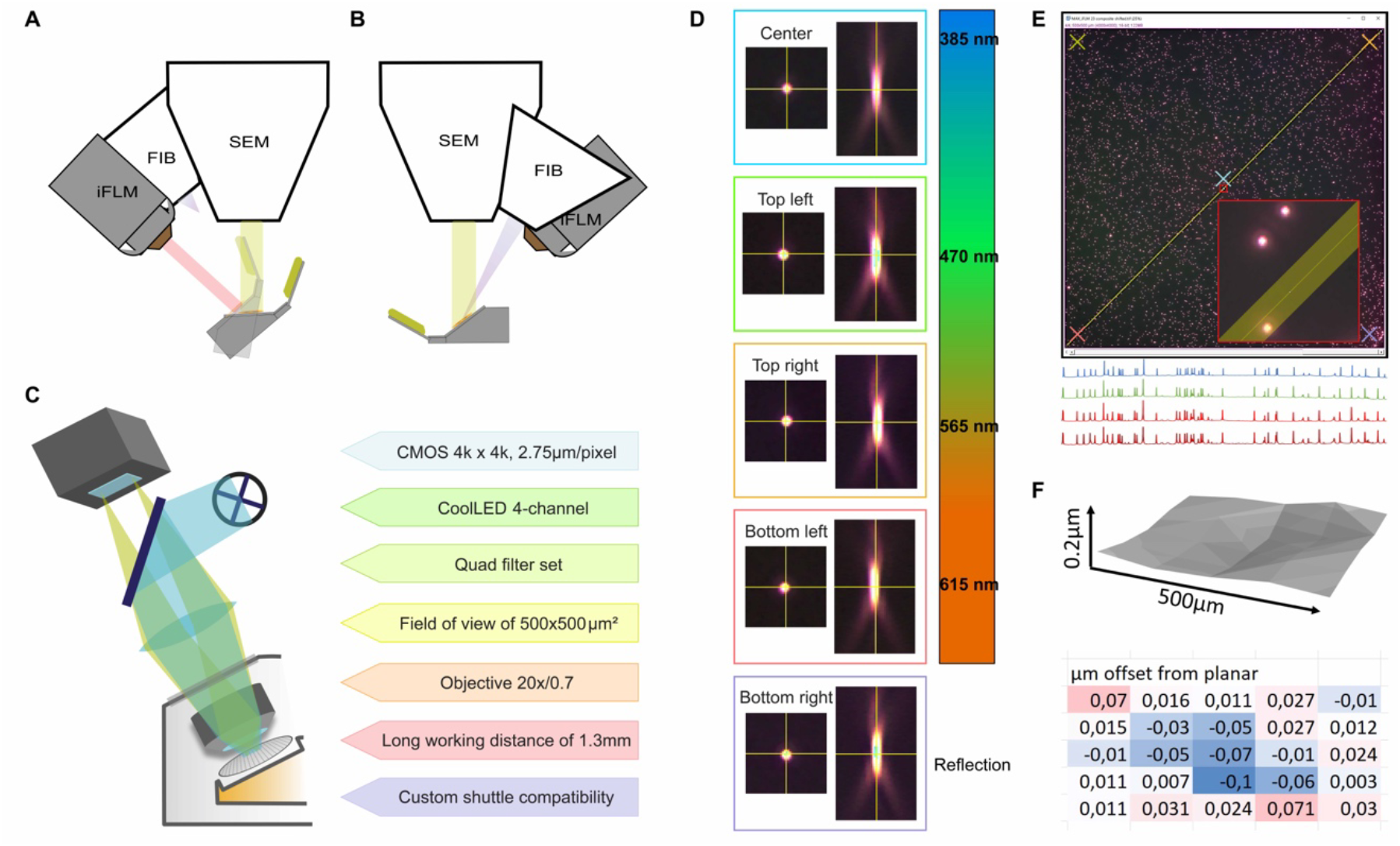
Integrated fluorescence light microscope (iFLM) in a cryo-FIB-SEM system. A schematic diagram of the in-chamber overview of the iFLM relative to the FIB and SEM columns, and the stage/shuttle position in the cryo-FIB-SEM system, from the front (A) and from the back (B). The stage moves laterally and tilts to the position where the fluorescence imaging angle is perpendicular to the sample surface under the iFLM module. When iFLM imaging is complete, the stage returns to the SEM imaging position where the viewing angle is perpendicular to the grid surface and milling position where the FIB ion-beam is positioned at a shallow milling angle. (C-F). iFLM resolution performance in fluorescence and reflected light microscopy modes. (C) An overview illustration of the iFLM features. (D) Chromatic aberrations demonstrated by the XY and XZ central slices of the point spread function (PSF) of four fluorescence and reflection channels via the Quad filter set of iFLM. (E) Flatness of the field of view under all four fluorescence channels demonstrated by the brightness of the beads in contrast to the background along the diagonal line of 700 *μ*m. (F) Curvature of the field of view under all 4 channels demonstrated by quantification of the z-offset of Tetraspec beads (200 nm in diameter) in 25 sub-fields of 100 x 100 *μ*m^2^.

**Figure 2.**
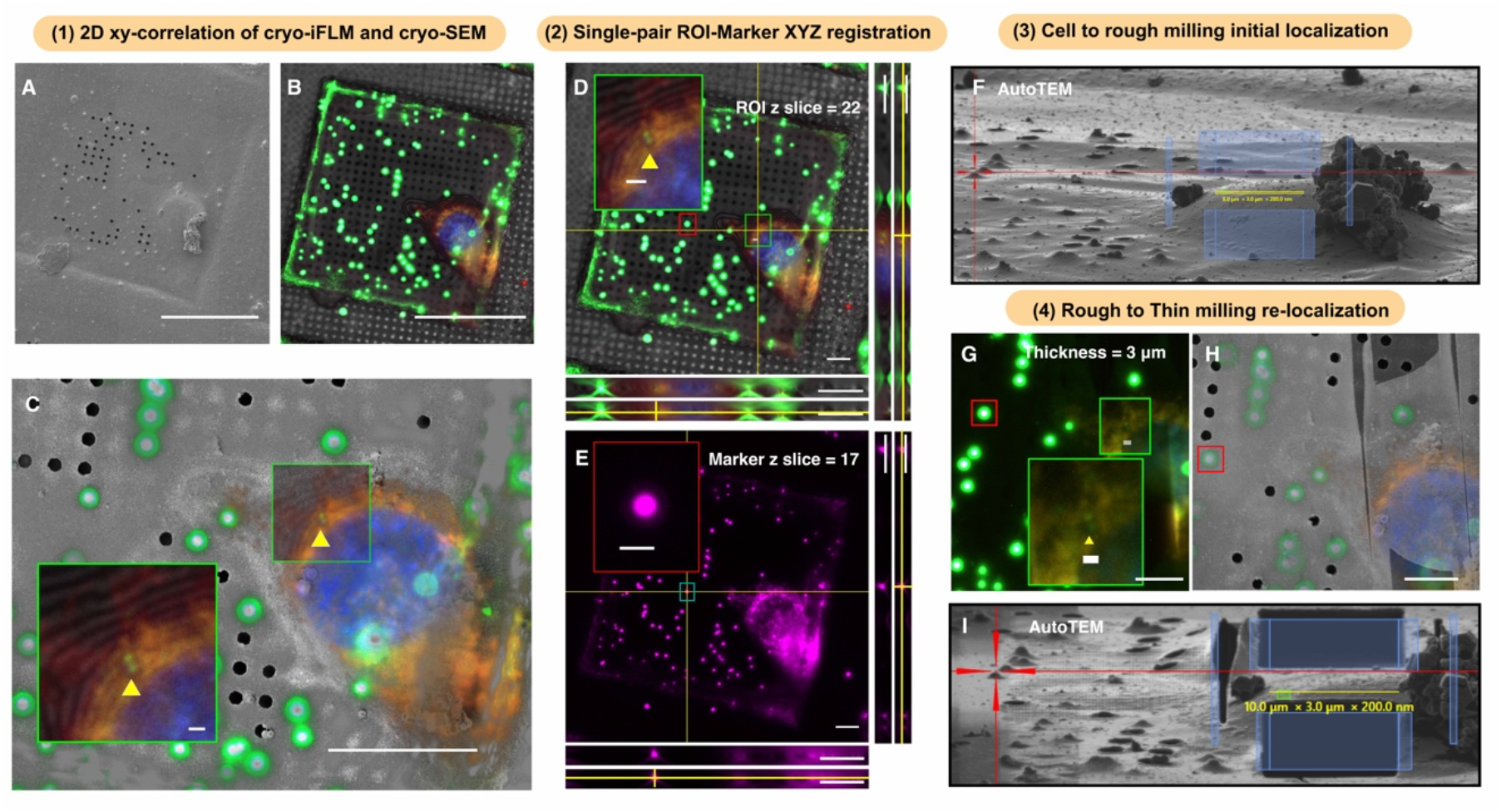
3D cryo-multi-modal correlation workflow in an iFLM-FIB-SEM system. iFLM guided FIB-milling procedure via the single pair ROI-Marker 3D correlation in a typical cryo-FIB milling session. (1) The registration of SEM (A) and iFLM (B) of a HeLa cell with lipid droplets (green), and mitochondria (red), and nucleus (blue) to show the XY location information of the ROI (single lipid droplet, yellow arrowhead). (2) The transformation of 3D Z positioning is achieved via the Z plane difference between the ROI (yellow arrowhead, D, Z slice of 22) and the marker (1 *μ*m bead, red box, em 605 nm, E, Z slice of 17). (3) The transformation of lateral information and Z-position from SEM/iFLM to the FIB via the milling angle on the targeted cell to set the initial location for rough milling. (4) A second round of registration and transformation between the same ROI-Marker in XYZ is performed at the end of the rough milling (thickness of 3 *μ*m). The automated milling process was not interrupted to produce ∼200 nm lamellae. The entire procedure is integrated in the AutoTEM automated milling workflow (AutoTEM 2.4.2). The screenshots of FIB correlative localization in AutoTEM (F and I). Scale bars of 50 *μ*m in A and B, 20 *μ*m in C (2 *μ*m in the zoom-in inset), and 10 *μ*m in D, E, G, H.

**Figure 3.**
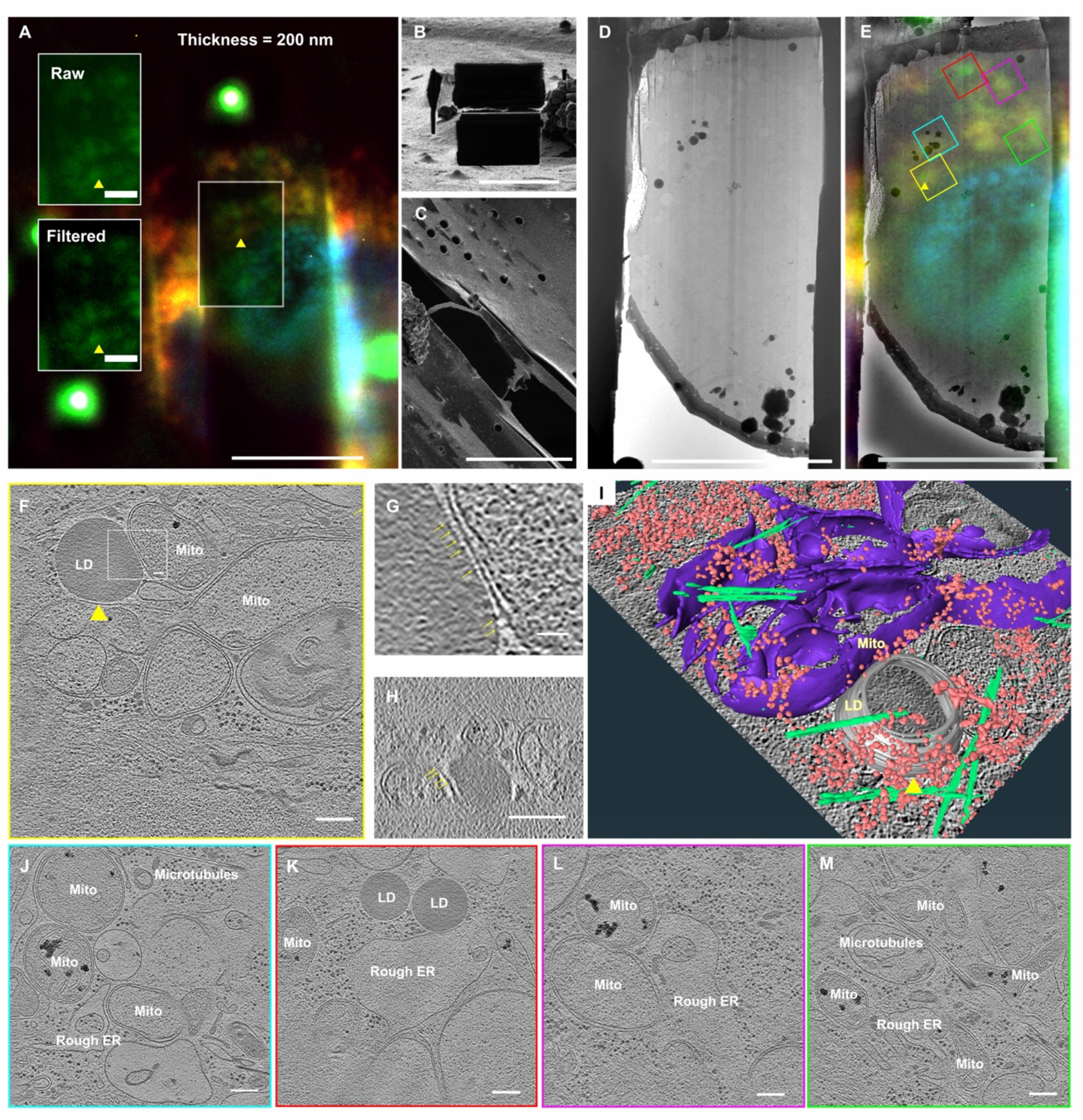
iFLM-guided FIB-milling and cryo-ET of HeLa cells to define LD-mitochondrial contact sites. The final 200 nm-thick lamella was generated by the single-pair ROI-Marker registration and transformation, imaged under the iFLM (A, Raw and deconvoluted views of same ROI (yellow arrowhead) and FIB (B viewed from the milling angle of 9°, stage rotated for sideview in C). (D-E) Cryo-EM image and overlayed iFLM image on the cryo-EM image of the 200 nm lamella. Note: preservation of the ROI for cryo-ET tilt series collection. The TEM images of the lamella (D-E) were oriented to align with the SEM/iFLM views. The orientation of the colored boxes matched with the actual TEM acquisition of the lamella’s orientation on the microscope stage. (F-M) Reconstructed tomogram XY slices of target sites (boxed regions) including the ROI of lipid-droplet (LD)-mitochondrial (Mito) contact sites (yellow boxed, yellow arrowhead in E-F, the enlarged view of the white boxed region in F where the LD connects with the mitochondria via protein-like density mediation shown in both XY view (G) and XZ view (H), 3D segmentation shown in (I) where microtubules (green), mitochondria (purple), ribosomes (pink), and ROI lipid droplet (yellow arrowhead, grey) were delineated), organization of mitochondria, microtubules, rough ER in the cellular environment (J-M, corresponding color boxed region in (E). Scale bars of 10 *μ*m in A-E, 200 nm in F, H, J, K, L, M, and 20 nm in G.

**Figure 4.**
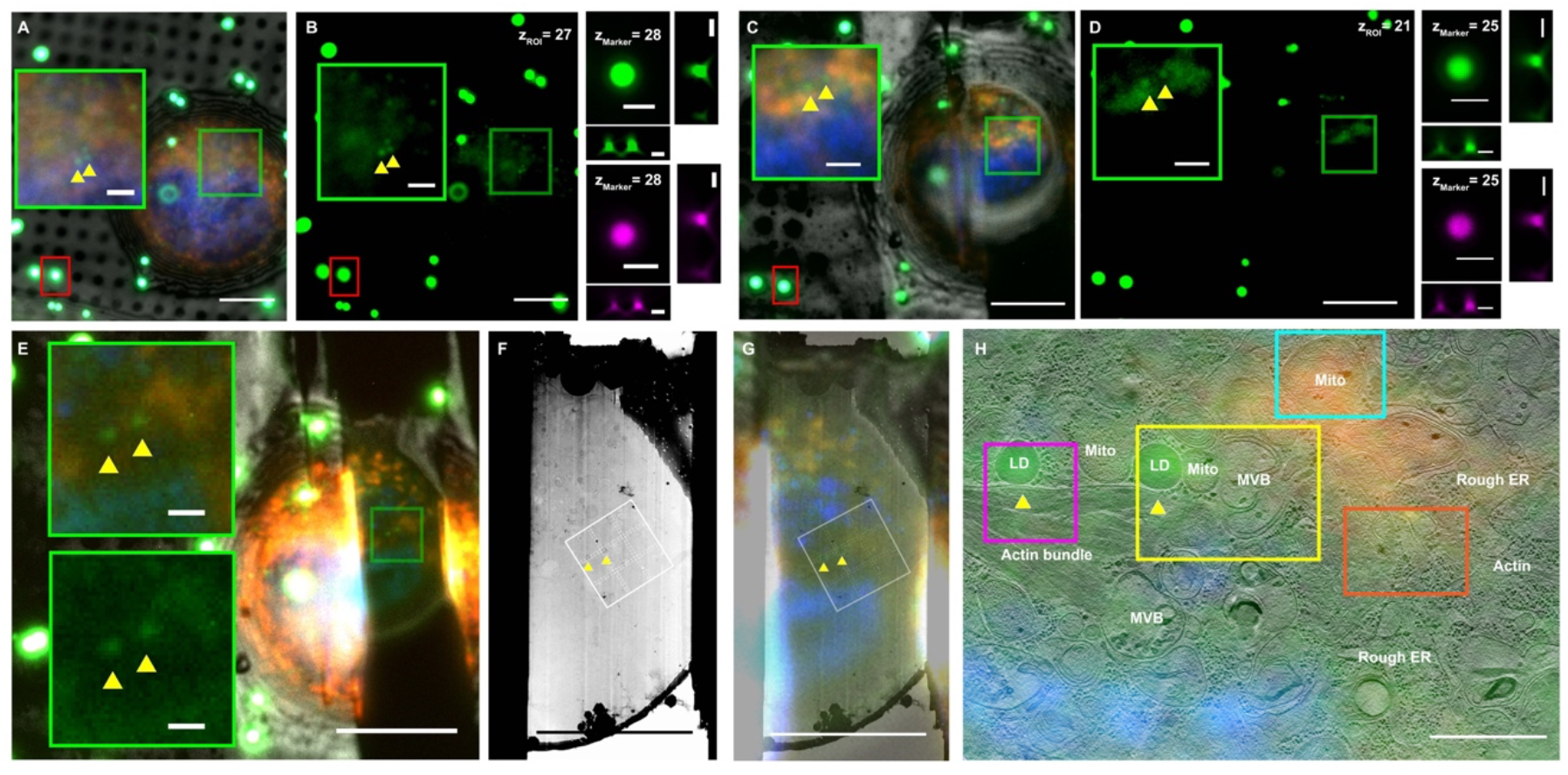
iFLM-guided FIB-milling and montage-cryo-ET (MPACT) of HeLa cells to study lipid droplet (LD)-mitochondrial contact sites. (A-B) 3D targeting of ROI for two adjacent LD and mitochondria. iFLM Z-stack plane differences in the ROI (Z-slice = 27 in the iFLM stack of the un-milled cell of interest, green box) and marker (Z-slice = 28 in the iFLM stack, red box) are presented. (C-D) A second round of single-pair ROI-Marker registration and transformation after rough milling of the lamella to 3 *μ*m thickness. The Z-positioning was updated via the z-plane differences of the ROI (Z-slice = 25 in the iFLM stack of the rough-milled lamella) and marker (Z-slice = 25 in the iFLM stack). After the second round of correlation, the automated milling process was not interrupted to produce ∼200 nm lamellae (E). (F-G) Cryo-EM image and overlayed iFLM image on the cryo-EM image of the 200 nm lamella. Note: preservation of the ROI for cryo-ET tilt series collection. The TEM images of the lamella (F-G) were oriented to align with the SEM/iFLM views. The orientation of the white 3x3 box (F-G) matched with the 3x3 cryo-ET montage acquisition via MPACT of the lamella’s orientation on the microscope stage. (H) Superimposition of the iFLM image and the reconstructed 3x3 cryo-ET montage. Colored boxed regions in (H) highlight subareas of note in the larger montage cryo-ET view. Scale bars of 10 *μ*m in A-G, 2 *μ*m in XZ and YZ views of the same marker beads before the milling (B) and after the rough milling (D), and insets in E, 1 *μ*m in H.

## Results

The principal design of the integrated FLM (iFLM) for the Aquilos 2 cryo-FIB-SEM system was for the FLM objective and the SEM-FIB probe-forming lens to maintain suitable working distances (WD) to support both imaging modalities and effortless region of interest (ROI) relocation between the two imaging positions. This iFLM arrangement is not exclusive to the Aquilos 2 system, and can be incorporated onto other commercialized cryo-FIB-SEM instruments such as the system published by de Marco et al (12). In the Aquilos 2 instrument, we placed the fluorescence light microscope (FLM) objective 1.3 mm away from the sample and parallel to the FIB column for recording images of the sample (Fig. 1A, B). The sample and stage can be reoriented to an ion perpendicular position by rotation of the stage and can also be repositioned to an electron perpendicular location by rotation of the stage by 180° and the application of tilt. This design supported free movement between all three imaging modalities. The iFLM module and its placement in the instrument had several advantages: 1) retention of a perpendicular (90°) incidence point to the sample surface allowed for the application of an affine transformation for correlation between the FLM and SEM, and the FLM and FIB; 2) a working distance (WD) of 1.3 mm for the FLM, compared to 1 mm (14); and 3) a piezoelectric positioner for focusing the FLM.

We selected, installed, and performance tested a 20x Zeiss Epiplan-Apochromat objective lens with an NA of 0.7; a field of view of 500 x 500 *μm*^2^ per frame, with an unbinned image pixel size of 0.125 *μm* at 20x magnification; and high-vacuum compatibility (Fig. 1C). To minimize pixel shifts between individual filter sets (Fig. 1D) and the time required to switch filter cubes, a multiband filter set (SI Appendix Fig. 1) was used for simultaneous and sequential imaging of multiple fluorophores, this was coupled with a CoolLED light source and room-temperature 4K x 4K CMOS camera (Fig. 1C). Use of a standard CMOS camera eliminated possible LM image blurring from internal vibration and permitted larger pixel densities not typically found with sCMOS. This additionally allowed for digital magnification changes between low and high magnification. This also helped with the navigation of large samples whilst not compromising high-resolution information. In addition to four fluorescence channels, bright-field imaging was supported by reflected light illumination, which was advantageous for surface-level imaging of the sample. To validate the performance of the optical microscope, we measured point spread functions (PSFs) for 200 nm Tetraspec fluorescent beads. The full width half maximum (FWHM) of the widefield PSF was 410 and 1600 nm for the XY and Z directions, respectively (470/510 nm channel). We quantified the field curvature aberration introduced by the LM lens and found insignificant variation to the planar offset (0.2*μm*, Fig. 1E, F). Raw iFLM fluorescence image stacks were further processed using standard deconvolution algorithms for improved lateral information in the XY plane (SI Appendix Fig. 2) to facilitate ROI identification and image contrast enhancement.

To correlate position and image data by the iFLM, SEM, and FIB, we developed and used a single-pair ROI-Fiducial-Marker registration strategy (SI Appendix Fig. 3 A-C). A 2D affine transformation was applied to the SEM and FLM images (SI Appendix Fig. 3 A, B), transferring the ROI coordinates from iFLM to SEM in the XY plane. Using a linear transformation, SEM images that contained the ROI coordinate information were projected onto FIB images acquired at the milling angles where the Y coordinate represented both translation in Y in the 2D XY plane and the Z height information in the 3D projection (SI Appendix Fig 3. C). Instead of directly using ROI coordinates for transformation, a fiducial marker reference point such as a FluoSphere, grid bar corner, or ice particle, was identified by both SEM and FIB. Therefore, the ROI could be projected onto the FIB view via its relative position to the fiducial marker. The actual z-location of the ROI was represented in the Y direction of the projected view (SI Appendix Fig. 3 C). This approach did not superimpose FLM-SEM to FIB for coordinate prediction and transformation, which simplified the 3D correlation process. The localization precision in this step was determined by the pixel size of the imaging modalities, and the manual selection of the ROI and fiducial marker positions. We then tested the correlation approach using 1 *μm* Fluospheres (ex/em 585/605 nm) at both room and cryogenic temperatures. We measured the relocation precision of the transformations by assigning one FluoSphere as an ROI (e.g., ground truth) and another as fiducial marker (SI Appendix Fig. 3 D). The relocation precision of the FluoSpheres at either RT or cryogenic temperatures were within 100 nm in X (xy, lateral) and 150 nm in Y (z-positioning). The target precision was independent of the milling angles used (SI Appendix Fig. 3 E). We also examined the use of milled reference patterns, such as an ‘L’ (SI Appendix Fig. 4 A), as fiducial markers because they could be identified by both SEM and FIB imaging modes. Relocation precision of milled fiducial markers imaged under reflection mode was comparable to FluoSphere-based registration under the FLM mode (SI Appendix Fig. 4 B). It is well known that cryo-milling of biological samples introduces internal stress within lamellae and subsequent deformation that could result in changes to the Z-height of the ROI. In addition, cryo-sample stages on Aquilos 2 systems may encounter mechanical drift that may shift the ROI in the XY direction. We monitored mechanical drift during the milling process by measuring the XY movement of lamellae. We selected distinct trackable features, including FluoSpheres and small chunks of surface ice near lamellae milling positions. The area was divided into two zones based upon the radial distance from the center of the target lamellae (interior: 16 *μm* diameter circle and exterior: beyond the 16 *μm* diameter circle). Shifts between 50 and 400 nm were observed in both zones during FIB-milling (SI Appendix Fig. 4). To mill the ROI with high accuracy, we implemented the single-pair ROI-Marker correlation in two steps (Fig. 2) and developed a dedicated workflow to perform an on-the-fly correlative milling procedure. Furthermore, to streamline this procedure and couple it with automated milling, we fully integrated the correlation steps in the AutoTEM cryo software program (v2.4, ThermoFisher Scientific). We were able to obtain 200-250 nm thick lamellae with preservation of the cellular ROI of 500 nm to 1 *μm* in size.

To benchmark this workflow, we targeted lipid droplets (LD) introduced to HeLa cells because of their distinct spherical profile for ROI identification and their biological significance in lipid metabolism and energy homeostasis by mitochondria (Fig. 2 and 3, LD noted by yellow arrowhead). In the first round of single-pair ROI-fiducial registration, FLM image stacks were collected that included the ROI within the cell and one nearby fiducial marker (1 *μm* Fluosphere bead) for assignment of XY/Z pixels (Fig. 2 (1)-(2)). AutoTEM-guided FIB-milling produced a ∼1.5 *μm* thick initial lamella that was not impacted by sample drift during the rough milling process (SI Appendix Fig. 4). In the second round of FLM imaging and registration, fiducial markers for the same ROI were selected and used. Based on the second round of correlation, the milling pattern was adjusted to guide thinning to produce a final 200 nm thick lamella that contained the ROI (Fig. 2(3)-(4)). The presence of the LD and mitochondrial targets preserved within the lamella was confirmed by iFLM inspection and cryo-ET data collection (Fig. 3A-E, ROI noted by yellow arrowheads). We observed densities associated with tethering structures between the LD and nearby mitochondria (Fig. 3F-I, SI Appendix Fig. 5A, yellow arrows). In the same lamella, we identified additional LD-endoplasmic reticulum (ER) contact sites, microtubules, and ribosomes (Fig. 3J-M).

To further validate iFLM-guided FIB-milling combined with large field of view tomography data collection by montage parallel array cryo-tomography (MPACT) (16), we investigated long-range cytoskeleton organization bounding LD-mitochondria rich regions in HeLa cells (Fig. 4). The target ROI (Fig. 4A-H, yellow arrowheads) remained centered in the final 200 nm thick lamella (Fig. 4E-H). Fully stitched montage tomographic reconstructions (3.7 x 5.4 *μm*) revealed diverse organizations of actin filaments that included loosely packed arrangements (SI Appendix Fig. 5 B-D), and densely bundled stress fibers (SI Appendix Fig. 5 E-G). Some stress fiber rich regions maintained higher order hexagonal-packing (SI Appendix Fig. 5 F-G). We measured differences in filament length and tortuosity between actin filaments (SI Appendix Fig. 6). This 3D correlative MPACT workflow enabled us to capture diverse cytoskeleton elements around LD-mitochondrial contact sites within their native environment to gain a more comprehensive 3D structural perspective.

## Discussion

In summary, we developed and present here a 3D correlative cryo-iFLM-FIB milling workflow using a commercially available system (Aquilos 2) and software programs (MAPS and AutoTEM cryo) to precisely target ROI of 0.5 to 1 *μm* in size within cryogenically preserved mammalian cells. Recently, a coincident three-beam cryo-FLM-FIB-SEM system was reported where FLM imaging could be performed during milling, supporting real-time monitoring (14). However, this configuration required the redesign of the cryogenic sample holder to allow the sample to be imaged from three sides simultaneously by FLM, SEM, and FIB. Consequently, the range within which the sample shuttle could move was limited to 28° of freedom in rotation at the coincidence point. The design and configuration of the iFLM in the Aquilos 2 high-vacuum FIB-SEM chamber ensured ample space necessary for the shuttle movements up to 54° of freedom in rotation range, perpendicular incidence point imaging on the sample surface in iFLM and SEM, and compatibility with the existing stage components. The space and stage movement freedom also permits operation of the Easylift needle for cryo-FIB Lift-out, which has become increasingly valuable for *in-situ* correlative cryo-iFLM-FIB milling workflows required for ‘bulk’ specimens such as tissue, organoids, and organisms.

Using the iFLM Aquilos 2 system, we demonstrated the accuracy of single ROI-Fiducial-Marker coordinate registration and on-the-fly relocation for 3D correlation under cryogenic-conditions. Linear transformations, such as an affine transformation, have been proven and preferred for 2D coordinate transfer for cryo-CLEM applications (5, 7, 18). Previous studies using a 3D rigid body transformation (7) demonstrated successful targeted cryo-FIB milling of cells. While elegant, this approach required a set of well-spread fiducials that need to be unambiguously identifiable near the target region. We established an efficient and simple method to register and transform coordinates from iFLM to FIB. The iFLM was able to provide real-time fluorescent signal spatial information after the grid was loaded into the cryo-FIB-SEM chamber. Using the locations provided by iFLM signals, we were able to apply a linear transformation and relative coordinate-based geometric alignment to accurately predict the ROI in both XY and Z within mammalian cells, via a single pair of ROI-Fiducial-Markers. We found this method not only suitable for fiducial-based registration, but applicable to sharp corners of milled reference patterns that serve as the Fiducial Marker. Additionally, it did not require the LM and FIB images to be superimposed, which greatly simplified the correlation process. To compensate for the movement of the lamella during the milling, possibly due to interactions with the ion beam, a second round of single-pair ROI-Fiducial-Marker registrations were performed on ∼1.5 *μm* thick lamella chunk after rough milling. This resulted in an overall localization precision of ∼300 nm or less.

Another common challenge with multi-fluorophore labeling in both conventional and cryo-FLM experiments is fluorescence signal bleed-through. Switching between single-band optical filter sets has been considered the gold standard for minimizing bleed-through background noise (SI Appendix Fig. 1 Bottom panel). However, this approach may introduce pixel shifts between multiple single-color images, and prolonged imaging times, which could make capturing dynamic processes challenging (17). In addition, there is an increasing demand for high-speed multicolor imaging approaches, which necessitates the use of alternatives to single-band filters/cubes without sacrificing image fidelity. As a result, a full multiband quad-band beamsplitter filter set coupled with a LED light source was chosen for the iFLM-Aquilos 2. Fluorescence signal from the FluoSpheres (ex/em 585/608 nm) was present in all channels and, in some cases, overpowered signals of interest, particularly, BODIPY 493/503 and MitoTrackerRed FM (SI Appendix Fig. 1 Top panel). FluoSpheres have significantly stronger absorption and emission profiles with extended tails when compared with many fluorescent dyes or markers of interest. To circumvent this problem, one may select FluoSpheres or other markers with different excitation/emission profiles that do not overlap or interfere with the fluorescence character of the labeled object of interest.

In conclusion, the 3D correlative iFLM-SEM-FIB workflow and registration strategy presented here provides a robust and efficient, automated correlation scheme with high-precision to produce thin cryo-lamellae of ∼200 nm with associated targeting errors of ∼300 nm. By fully embedding the correlation workflow within the AutoTEM cryo automated milling regime, FLM-targeted high-throughput milling is achievable. Coupled with the previously developed montage parallel array cryo-ET (MPACT) (16), precise 3D iFLM-guided targeted-cryo-ET supported studies of complex actin networks near LD-mitochondrial contact regions in the context of a native cellular environment. Future developments could include the expansion of correlative imaging workflows with the iFLM-Aquilos 2 system, such as alternative correlation algorithms, correlation with external FLM instruments, and additional integration with 3D cryo-lift-out applications.

## Materials and Methods

### Cell Culture, Seeding, Live-Cell Labeling on TEM Grids

HeLa cells (ATCC CCL-2, ATCC, Manassas, VA, USA) were cultured and maintained in supplemented DMEM complete medium as reported previously. HeLa cells were trypsinized and seeded at a density of 0.5 – 0.7 X 10^5^ cells/mL on carbon-coated gold Quantifoil grid (200 mesh, R 2/1, holey SiO_2_ film; Quantifoil Micro Tools GmbH, Großlöbichau, Germany) for 8 to 12 hr to allow for cell attachment and growth. To visualize mitochondria, nuclear DNA, and lipid droplets, grids seeded with HeLa cells were washed with warm 1x PBS, stained with MitoTrackerRed FM (M22425, ThermoFisher Scientific, 200 nM, 20 min at 37°C and 5 % CO^2^), washed twice with warm 1x PBS, subsequently stained with Hoechst-33342 (H3570, ThermoFisher Scientific, 1:1000 dilution, 20 min at 37 °C and 5 % CO^2^), washed twice with warm 1x PBS, followed by a final staining with BODIPY 493/503 (D3922, ThermoFisher Scientific, 3.8 *μ*M, 15 min at 37 °C and 5 % CO^2^). The stained cells were washed twice with warm 1x PBS prior to plunge-freezing.

### Vitrification

The grids were plunge-frozen using the Leica EM GP plunger (Leica Microsystems, Germany). The plunger was set to 25 °C to 30 °C, 99% humidity and blot times between 12 and 15 s. An additional 5∼10 % glycerol incubation of the grid prior to the freezing was performed to ensure proper vitrification as reported previously (19). Afterwards, 4 *μ*l of 1 μm FluoSpheres (1 to 800 dilution, 585/608 nm, ThermoFisher Scientific, F13083), were applied to the grid in the humidity chamber of the Leica EM GP plunger prior to the vitrification step. The plunge-frozen grids were then clipped and stored in cryo-grid boxes under liquid nitrogen.

### Integrated Fluorescence Light Microscope (iFLM)-Aquilos 2 Setup and Data Acquisition

The commercial iFLM-Aquilos 2 equipped with a 20x 0.7 NA, 1.3mm WD objective lens (Zeiss Epiplan-Apochromat) for fluorescence imaging was introduced to the 3D correlative workflow. An LED illumination source was used, a multi-band-pass dichroic (Semrock LED-DA/FI/TR/Cy5-B-000 (Quadband)) and customer filter exchanger which allowed for both fluorescence and reflected light imaging. The light source is a CoolLED, 4 channels (365 nm, 450 nm, 550 nm, 635 nm). Images were captured on a RT Basler ace 2 (2A4504-5gmPRO; Sony IMX541 CMOS sensor) 4K x 4K detector. To position the iFLM closer to the electron and ion beam columns, the ICE detector was relocated. Fluorescence imaging tasks were controlled by a custom python GUI using AutoScript to control the microscope. These included direct movements between imaging modalities (SEM to FLM, FLM to SEM, FIB to FLM, and FLM to FIB), sample focusing, Z-stack image collection, and multi-channel fluorescence and reflection imaging.

All iFLM data sets were recorded using the multiband quad-band beamsplitter filter set (Fluorophores selected: DAPI/385 nm, FITC/470 nm, TRITC/565 nm, CY5/625 nm) at bin 1x on the CCD camera. The intensity of the light source was set to 35% relative to the full power. The exposure time used varied from 5 msec to 500 msec, depending on fluorescence signal. We did not observe prominent sample damage, validated via cryo-TEM, when the intensity power under bright field imaging was as high as 80 % with a long exposure time of 1000 msec.

### Image Processing of Raw Fluorescence Image Stacks from the iFLM

The iFLM image stacks were deconvolved using standard Richardson-Lucy (RL method. (20)) implemented in DeconvolutionLab2 (21) in Fiji (22). The point spread function (PSF) used was directly obtained by imaging 200 nm Tetraspeck beads under the four fluorescence channels via the iFLM objective lens operating under cryogenic conditions. The optimal number of iterations was determined heuristically to stop the algorithm before convergence for each fluorescence channel. In this manner, artifacts were minimized resulting in a lateral resolution improvement in XY. The intensity profile in X was plotted in Fiji using *Plot Profile*.

### 3D Correlation Via Single-Pair ROI-Marker Registration In MAPS And AutoTEM cryo

The on-the-fly cryo-iFLM-SEM-FIB coordinate and stage position transformations implemented here were based on a linear transformation. The perpendicular incidence point of the beam to the sample surface in both iFLM and SEM imaging allowed for 2D rigid affine transformations to be used to transform the XY plane 2D locations from the iFLM (fluorescence or reflection mode) to the SEM view. In 2D plane projections, the SEM and FIB have the same coordinate information in the X direction. Therefore, the points identified in iFLM could be directly transferred from iFLM/SEM to FIB after the initial iFLM to SEM correlation.

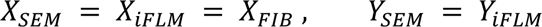

With a flat sample, 2D projection of an object via scaling transformations represents the 3D rotation of an object around the X- and Y-axis. The 2D linear transformation can be used for the rigid 3D transformation of a flat object. Thus, the Z information of a nearly flat ROI in iFLM, e.g., microspheres (diameter < 1 *μm*) could be represented by the 2D rigid transformed Y coordinate projection in the FIB. To correctly transform a non-flat 3D object from iFLM to FIB, e.g., biological samples, the Z height information of the object relative to the support surface (XY plane) needs to be incorporated. This yields

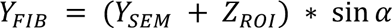

where α is the milling angle. It is implicitly understood that the *Y*_%’(_ coordinate represents both the 2D XY-plane and the 3D Z-height information related to the 2D projection of the ROI. In practice, we found that the relative positioning of ROI to a nearby “Fiducial Marker” object relative XY and a height difference in Z could serve as one unambiguous registration point. This offers an advantage over direct ROI coordinate correlation where multiple registration points (*n* ≥ 5) need to be well-spread for accurate transformation calculations. By identifying the single Fiducial Marker point, we could locate the ROI based on the relative position information.

Here, potential milling sites were identified based on fluorescent signals of interest, e.g., lipid-droplet puncta (green) or mitochondria (red) sites in HeLa cells. The FluoSphere (diameter of 1 *μm*) auto-fluoresced under both green and red channels. The ROI within a targeted cell was identified as an “ROI” at the focal plane (Z_ROI_) in the green channel. Correspondingly, one FluoSphere near a target cell within the same square was identified as “Fiducial Marker” at the focal plane (Z_Marker_) in the red or green channel to minimize the errors due to the chromatic shift between channels.

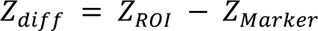

Then *Y_FIB_* that represents both the 2D plane in Y axis and Z height information in 3D is below:

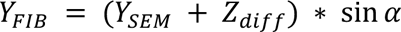

Functions that perform all calculations and correlations mentioned above, such as the 2D linear transformation between SEM and iFLM, relative Z height information, identification of single-pair ROI and Fiducial Marker, transformation between image coordinates to stage positions, are all embedded in MAPS 3.21 and AutoTEM cryo 2.4. Therefore, the 3D correlative workflow is coupled with the automatic milling process.

Two rounds of iFLM imaging were conducted in this workflow. Briefly, the clipped HeLa cell grids were loaded onto the iFLM-Aquilos 2 operated under standard cryogenic-condition (-195 °C). The first round of iFLM imaging using the iFLM software was performed on unmilled HeLa cells after the sample was sputter-coated with Platinum (20 mA, 25 sec), and subsequently coated with organometallic Platinum using the in-chamber gas injection system (GIS, 75 sec with a measured deposition rate of 60 nm/sec). Optionally, a second Platinum sputter coating (20 mA, 7 sec) was performed to reduce sample charging effects. The fluorescence and reflection z-stack projections of 12 to 15 *μ*m were collected with a step increment of 350 nm. The .xml files for each iFLM image/stack that contained both image and stage coordinate transformation information were loaded into MAPS 3.21. A 2D linear transformation was performed between the SEM and iFLM stack at the correct focal plane in either the reflection or fluorescence channel, using the “Alignment” function in MAPS. The single-pair ROI-Fiducial Marker was identified in MAPS 3.21. This transformation and relative positioning was transferred and directly linked to the “Correlation” function within the “Preparation” step in AutoTEM cryo 2.4. The Marker (red cross in AutoTEM cryo) was first manually identified in the FIB view and the ROI (green box in AutoTEM cryo) was automatically located based on the calculation. The initial rough milling boxes were then placed around the ROI. Rough milling to generate lamellae of 1.5 *μm* in thickness was conducted at 0.3 nA. To compensate for lamella movement during milling and to refine relocation of the ROI, a second round of iFLM imaging for 3D correlative positioning was performed post rough milling. Similarly, the z-stacks of both fluorescence and reflection channels were collected at each milling site. The same ROI was identified on its focal plane (Z_ROI_). A nearby FluoSphere was then used as a Fiducial Marker, identified on its focal plane (Z_Marker_). The updated single-pair transformation was loaded into the AutoTEM cryo “Preparation” step where an updated Medium milling box positioning step could be conducted. The automated milling process was continued through to the “Polishing” step to obtain ∼200 nm lamella in AutoTEM cryo 2.4. Parameters in the “Polishing” step might need to be optimized based on the sample type, lamellae positions, grid square integrity (e.g., the presence of cracks). Prior to removal from the cryo-FIB-SEM, an optional additional iFLM imaging step could be performed to verify and confirm the preservation of ROI.

To use sharp corners as Fiducial-Markers, L-shaped rectangular milling patterns were placed and created using a current of 0.1 nA, after sputtering and GIS preparation. The milled fiducials were located 15∼20 *μm* away from targeted cells and ROIs on the support surface. The shuttle stage was rotated 180° relative to the milling stage position and tilted to 17°, to ensure a perpendicular ion-beam incidence to the sample surface. The corners of the milled L-shape were used as Fiducial-Markers during the first and second rounds of single-pair ROI-Fiducial-Marker registration and correlation steps, as described above.

### Alignment Accuracy Estimation

We validated correlation relocation accuracy for single-pair Fiducial-Marker-ROI registration at RT and cryogenic conditions, using FluoSpheres (diameter of 1 *μ*m) on gold Quantifoil grids (200 mesh, R 2/1, holey SiO_2_ film; Quantifoil Micro Tools GmbH, Großlöbichau, Germany). Briefly, the FluoSphere coated grids were sputtered and GIS coated using the same conditions as biological samples. FluoSphere pairs within one grid square were randomly selected. One served as the Fiducial Marker and the other as the ground-truth ROI whose position was predicted in the FIB view in after correlation. The deviation in the X and Y axes was calibrated as a Euclidean vector. The deviation in Y axis of the predicted ROI against the true ROI provided the alignment estimation of the Z height. The centroid of the FluoSphere ROI and Fiducial Marker was determined in MAPS 3.21 via the iFLM z-stack. The deviation measurements were done in Fiji using the FIB view and predicted ROI green box from AutoTEM cryo after the Marker registration.

### Tracking Movements of Lamellae and Neighboring Areas Under Cryogenic Conditions

The movements of the lamella, during the entire milling process were sequentially recorded via SEM with an interval of 30 sec (2kV, 25 pA, 3072x2048, dwell time of 1 *μ*s), using the iSPI tool within xTUI (32.0). No stage movement or drift corrections were performed or prevented. The SEM image stack was filtered in Fiji using a smoothing algorithm (kernel of 2) to improve the contrast. A series of MATLAB programs for Tracking and Motion Estimation were used to automatically recognize and follow the movement of features throughout the image stack. Briefly, on the first frame, the feature points for tracking were either automatically identified using MATLAB (R2020b and above, MathWorks Inc.)-based program *detectHarrisFeatures*, or manually seeded in Fiji and saved as pixel coordinates. The MATLAB program *PointTracker* was then applied to follow the movements of the feature points throughout the stack. The movement in pixels was recorded per point per frame and a tracing line was drawn to delineate the paths of the feature. The standard deviation (StD) of the coordinates was used to represent the how much movement occurred during the process. Bigger StD indicated larger movements. The global movement on the whole frame level was estimated via the MATLAB program *BlockMatcher*. The tracking procedures were developed into three MATLAB automated scripts, available upon request.

### Cryo-Electron Tomography and MPACT Acquisition

After 3D correlative cryo-iFLM-FIB-milling, the grids were imaged using a Titan Krios G4 with a cold Field Emission Gun (E-CFEG)-TEM (Thermo Fisher Scientific, Hillsboro, OR, USA) operated at 300 kV. Images were acquired on a SelectrisX-Falcon 4i direct electron detector (Thermo Fisher Scientific, Hillsboro, OR, USA) in EFTEM mode using a 10-eV slit. Images were captured at magnifications of 125x (1007 Å/pixel) for whole grid mosaic collection, 1950x (63.58 Å/pixel) for whole lamella overview, 6500x (19.26 Å/pixel) for intermediate magnification imaging for feature identification and tracking, and 26000x (4.727 Å/pixel) for data acquisition using SerialEM software (v.4.1). Full frame images of 4096 x 4096 pixels (counting mode at 2∼10 e-/px/s (eps) of dose rate over the sample) were collected and saved in the Electron Event Representation (EER) format. Tilt series were collected using a dose symmetric scheme with 3° increments, groups of 2 tilts, a nominal defocus of -8 *μm*, and a total dose of ∼80 *e*/Å^2^. Additional requirements for Montage Parallel Array Cryo-Tomography (MPACT) were 3x3 montage cryo-tilt series with 12 % overlaps in both X and Y, and a total dose of ∼50 *e*/Å^2^ per each tile tilt series were collected. The default spiral setting (A_final_ = 1.5, Period = 3, Turns = 50, Revolutions = 15) was used to spread the dose, as reported previously (16).

### Cryo-Tilt Series and MPACT Tomogram Reconstructions

All raw movie frames/fractions per acquisition (4096 x 4096, counting, EER format) were grouped into fractions to achieve 0.1 ∼ 0.3 *e*/Å^2^/fraction, brought to 8K super-resolution from a physical 4K grid for alignment and motion correction via MotionCor2. For regular cryo-ET, the motion-corrected frames were then sorted to generate new tilt series (pixel size of 4.727 Å^2^/*pixel*) that were aligned and reconstructed with AreTomo (23). The final tomograms were binned by 2 (final pixel size of 9.454 Å^2^/*pixel*). For MPACT tilt series, individual motion-corrected tile frames were first stitched per tile and then used to generate a stitched tilt series by running an automated pre-processing pipeline as reported previously (16). The stitched tilt series were binned by 2 (pixel size of 9.454 Å^2^/*pixel*), aligned via patch tracking and reconstructed using IMOD (24). Raw tomograms, both regular and montage cryo-ET, were processed using IsoNet (25) at pixel size of 18.908 Å^2^/*pixel*, to reduce the missing wedge effect and improve the signal-to-noise ratio.

### Fiber Tracing Analysis and Segmentation

In order to analyze actin organization, processed tomograms (binned 4X, pixel size of 18.908 Å^2^/*pixel*) were imported into Amira (ThermoFisher Scientific) for fiber tracing with the TraceX plugin (26). Briefly, Cylinder Correlation was performed (cylinder length of 80 nm, angular sample of 5°, outer cylinder diameter of 4 nm, mask cylinder of 5 nm, missing wedge correction on), followed by Trace Correlation Lines (direction coefficient of 0.15, minimal distance of 12 nm, search cone length of 80 nm with angle of 37°, and minimal step size of 10%, minimum seed correlation and minimum continuation quality were tomogram dependent). Fiber tracing results were checked with Spatial Graph View and non-actin features such as membranes were edited in the Filament module in Amira. The coordinates of all fibers were extracted, exported as xml files, and imported into the MATLAB Ultrastructural/computational toolbox (27). Visualization and quantitative parameter analyses including length, bend, cross-sectional circularity, filament angles, were conducted tomograms of the lamellae.

Cellular tomograms were segmented in Amira (ThermoFisher Scientific). The membranes (e.g., mitochondria) were generated automatically using a *Membrane Enhancement Filter (28)*, *Hysteresis Thresholding*, and *Filtering by Measurement* functions. The ribosomes and microtubules were segmented via the Deep Learning neural network tool, while the lipid droplets were traced manually in the Segmentation workroom.

## Acknowledgments

We are grateful for the TEM and SEM instrumentation support of Mr. Micky Woods and Mr. Tom Coomes from Thermo Fisher Scientific. We are grateful for the computational resources supplied through the SBGrid-supported (29). This work was supported in part by the University of Wisconsin, Madison, the Department of Biochemistry at the University of Wisconsin, Madison, the Morgridge Institute for Research, and public health service grants R01 GM114561 and U24 GM139168 to E.R.W. from the NIH. This work was supported in part by the U.S. Department of Energy, Office of Science, Office of Biological and Environmental Research under Award Numbers DE-SC0018409. We are grateful for the use of facilities and instrumentation at the Cryo-EM Research Center and the Midwest Center for Cryo-ET in the Department of Biochemistry at the University of Wisconsin, Madison.

## Supplementary Information Appendix

**Fig. S1.**
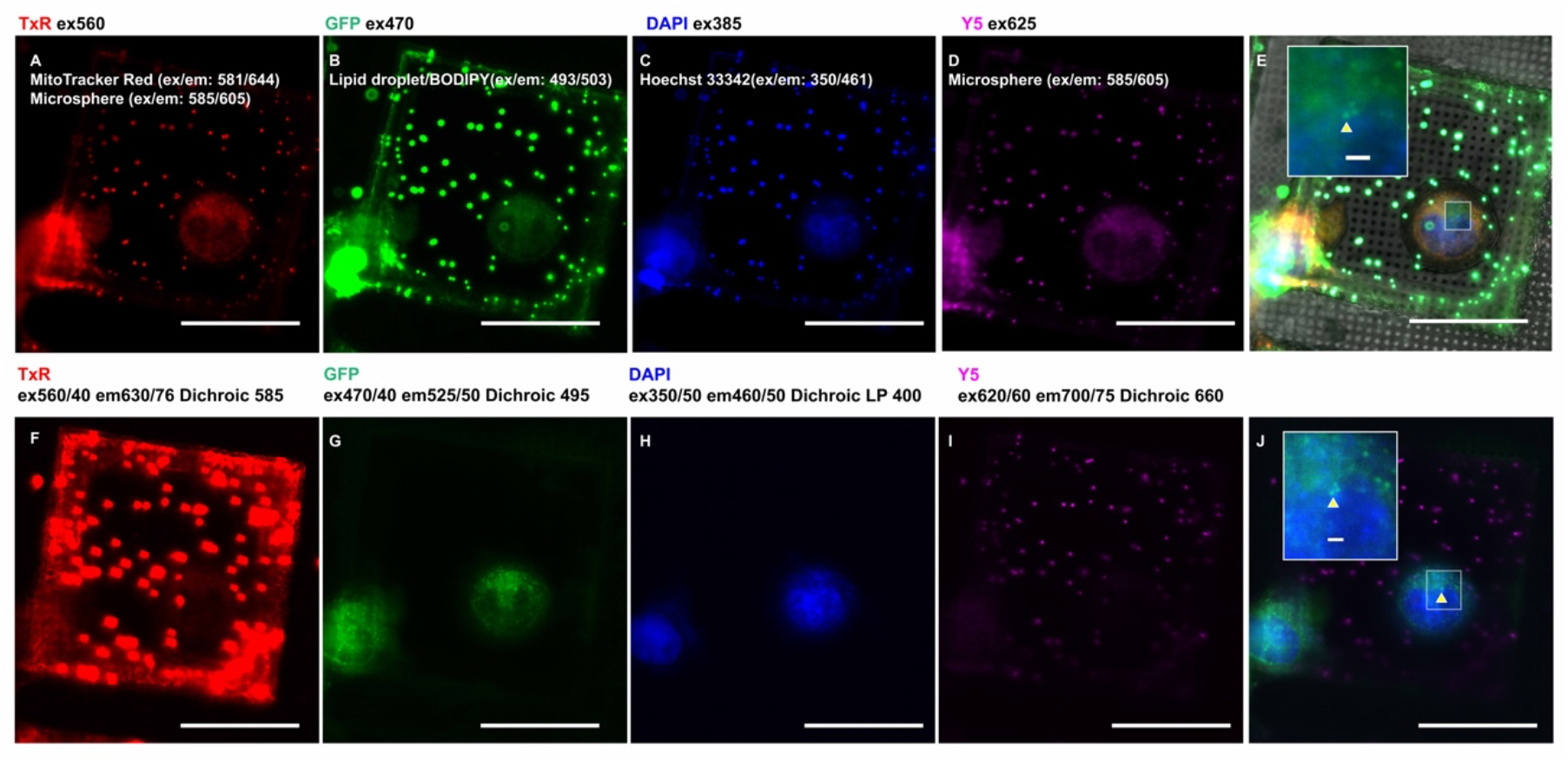
Comparison of transmittance performance of the multi-band pass filter and a single-band pass filter under cryogenic conditions. The cryogenic transmittance performance of multiple fluorescent signals was examined using the same plunge-frozen on-the-grid HeLa cells with labeled mitochondria (red, MitoTracker, ex/em 581/644), lipid droplets (green, BODIPY, ex/em 493/503), nucleus (blue, Hoechst, ex/em 350/461), and an addition of external 1 ***μ***m microspheres (ex/em 585/605) as registration markers. The top panel (A-E) shows the multi-band-pass filter performance of iFLM-Aquilos 2. The bottom panel (F-J) represents the single band-pass filter cubes of a Leica THUNDER cryo-CLEM system. Both systems are equipped with ceramic objective lenses that have similar numeric aperture values (NA of 0.7 in iFLM-Aquilos2 and NA of 0.9 in Lecia cryo-CLEM) and RT CMOS-based detectors, operating in the wide-field, with the samples under cryogenic conditions. Scale bars of 50 ***μ***m in A-J, 2 ***μ***m in the inset of (E) and (J). Yellow arrowheads in (E) and (J) note the ROI fluorescence imaging targets.

**Fig. S2.**
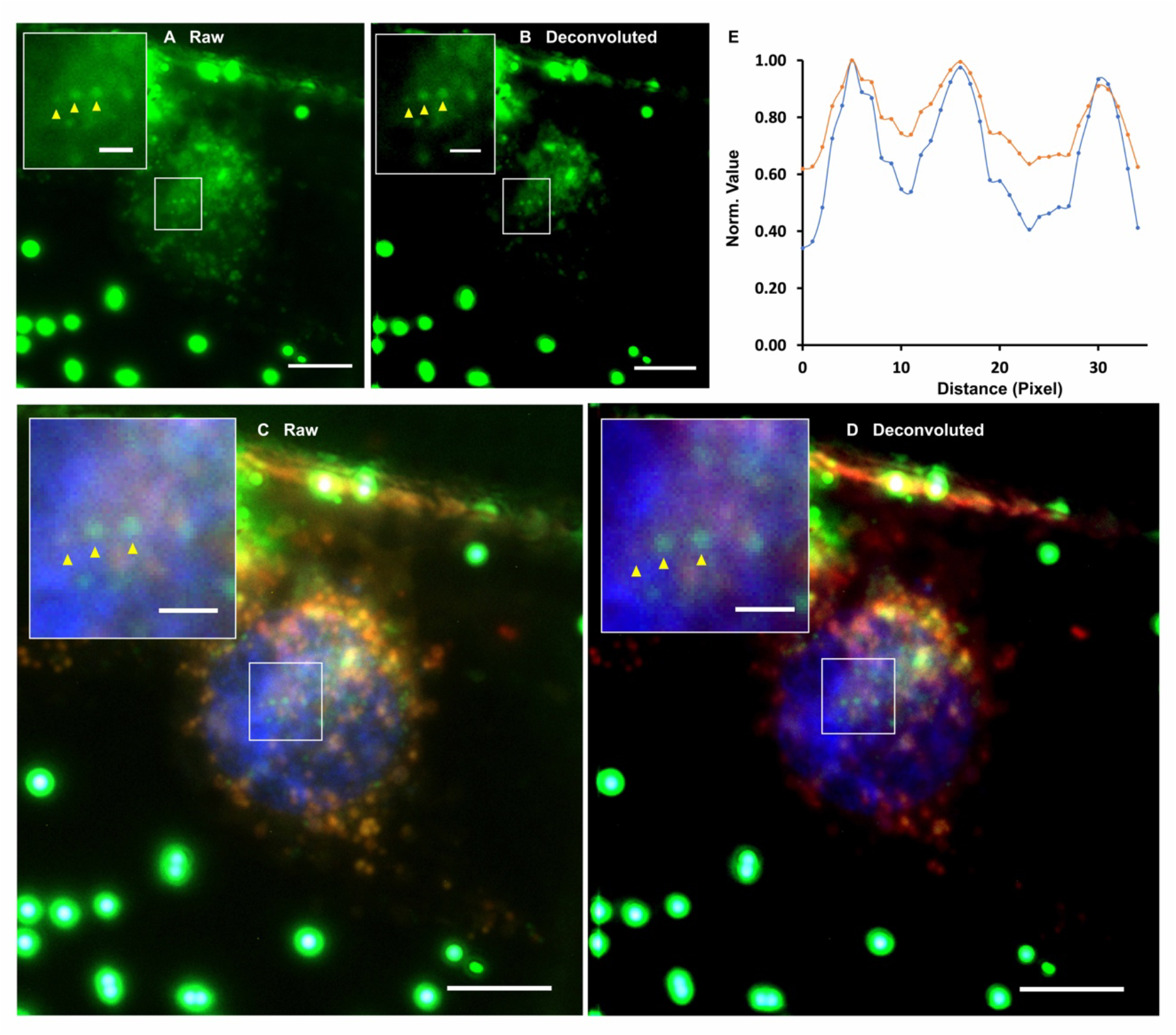
Improved contrast and image resolution of iFLM after deconvolution. (A) The three green ROI (yellow arrowheads) were targeted and shown in a single Z-plane of a raw iFLM image stack of a fluorescently labeled HeLa cell (red, MitoTracker, ex/em 581/644), lipid droplets (green, BODIPY, ex/em 493/503), nucleus (blue, Hoechst, ex/em 350/461). (B) The corresponding Z-plane of the PSF-deconvoluted iFLM image stack. Yellow arrowheads in (A) and (B) point out the ROI fluorescent targets. Scale bars are 50 ***μ***m in (A-B), 2 ***μ***m in the inset of (A-B). (C-D) The resultant merged multi-fluorescent image focal plane of raw (C) and PSF-deconvolved (D) iFLM image stack. (E) Normalized X-axis intensity plot file of the three ROI in the raw (orange line) and deconvolved (blue) image plane show undetectable pixel changes after the deconvolution processing with a clearly improved lateral PSFs of ROIs (yellow arrowheads in A-B). Scale bars of 50 ***μ***m in (C-D), 2 ***μ***m in the inset of (C-D).

**Fig. S3.**
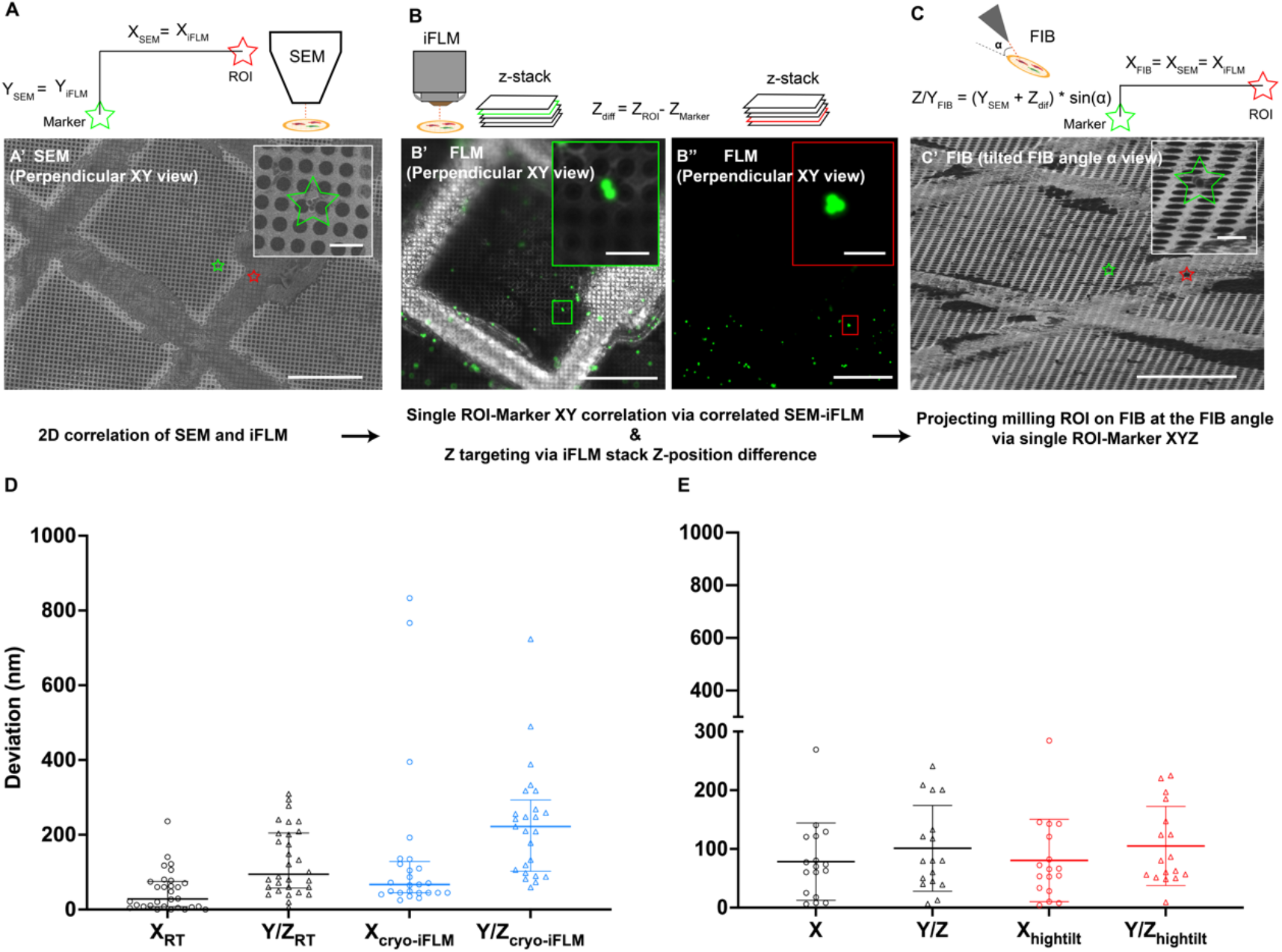
Coordinate-based two-point 3D correlation in an iFLM-FIB-SEM system. (A-C) Single-pair ROI-Marker registration of iFLM and SEM and its correlation projection on the FIB view to locate the ROI. The configuration of SEM and iFLM objectives allows for perpendicular projection of the sample on the shuttle where the lateral location in X is equivalent across iFLM, SEM, and FIB after correlation between SEM and iFLM (A, C). The Z-positioning takes into consideration the milling angle (***α***) and the true Z height difference between the ROI and Marker provided by the 3D iFLM Z-stack (B). The 3D ROI relocation in XYZ is achieved in the FIB view using the single ROI-Marker registration (C) where the Y coordinate represents the 3D Z-positioning information. (D) Targeting precision of the direct, unsupervised ROI localization for 1 ***μ***m microspheres (em 605) under RT (n = 30) and cryo-conditions (n = 25) in X and Y/Z. (E) Precision of the targeting ROI location under RT of the same 1 ***μ***m microspheres (n = 17) (em 605) in X and Y/Z in shallow milling angle (9°, black data points) and high milling angle (21°, red data points). A known 1 ***μ***m microsphere as the ROI to serve as the ground truth and a nearby 1 ***μ***m microsphere selected as the marker for registration in both (D) and (E).

**Fig S4.**
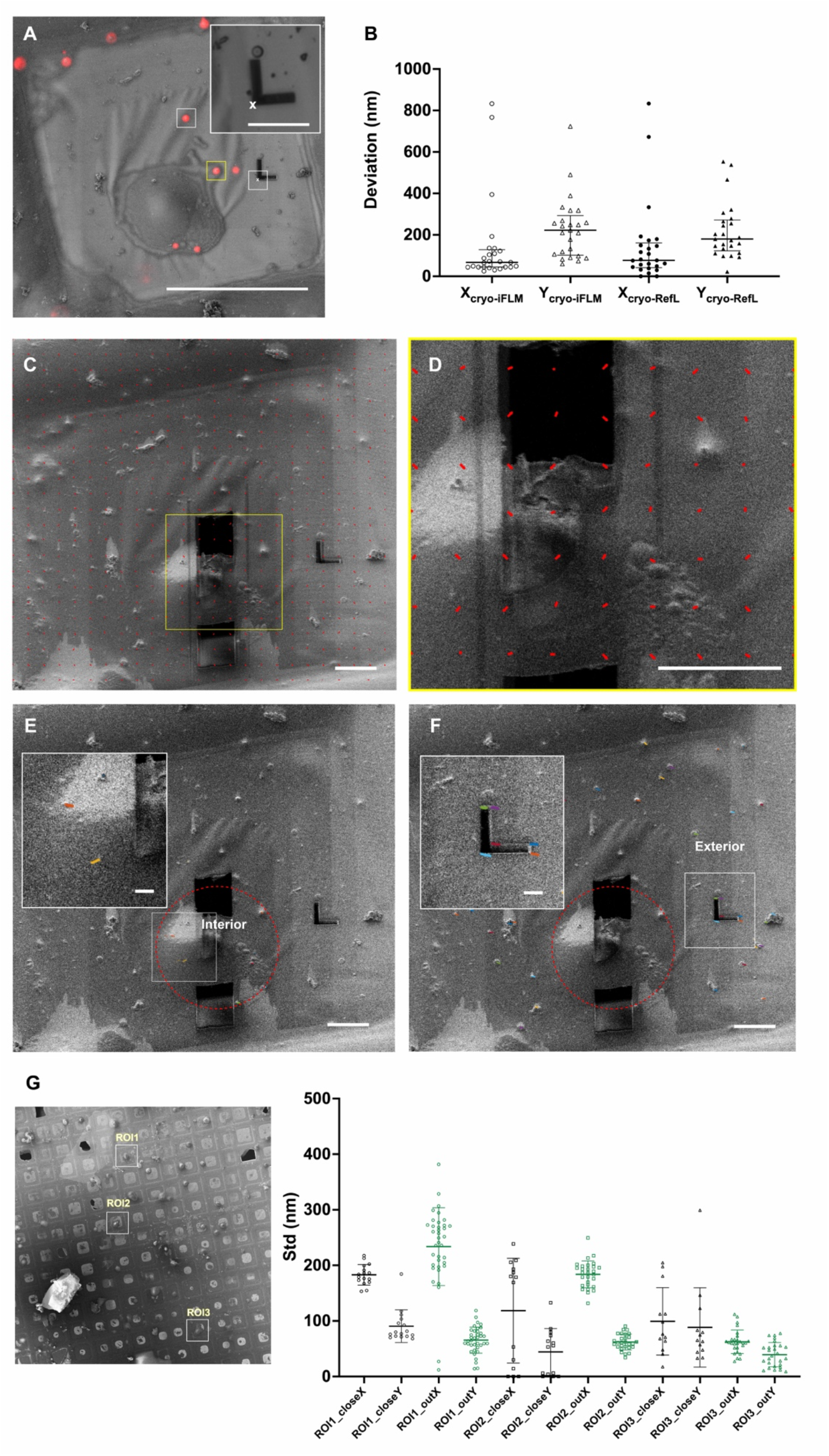
Marker registration strategies and the movement of fiducial markers during cryo-FIB milling. (A) A microsphere bead (white box) served as a fiducial marker point. The sharp corner of a L-shaped FIB-milled pattern (white box with an asterisk to indicate the selected corner) on the support film within the same grid square was an alternate fiducial marker for the single-pair ROI-marker registration/transformation (yellow box). (B) Targeting precision comparison of the same sets (n = 25) of fluorescent 1 ***μ***m microsphere as the ROI using either nearby 1 ***μ***m microspheres in the fluorescence channel or the sharp corner of a L-shaped milled pattern in the reflection mode under the cryogenic conditions. (C-D) Global movement of objects during milling (red arrows located in sub-regions with dimensions of 0.5 x 0.5 ***μ***m^2^). (E-F) Local movement of the objects during the lamella milling process (E, interior, 16 ***μ***m in radius, red circle), and the outer region (F, exterior). (G) Quantitative analysis (Right, G) of the movements of three frozen mammalian ROI milling sites on a representative grid (Left, G) via tracking the interior and exterior landmarks throughout the entire FIB-milling process. The movement is presented via the standard deviation (StD) of the coordinate in X and Y of the landmark. Larger StD indicates larger movements of the landmark.

**Fig. S5.**
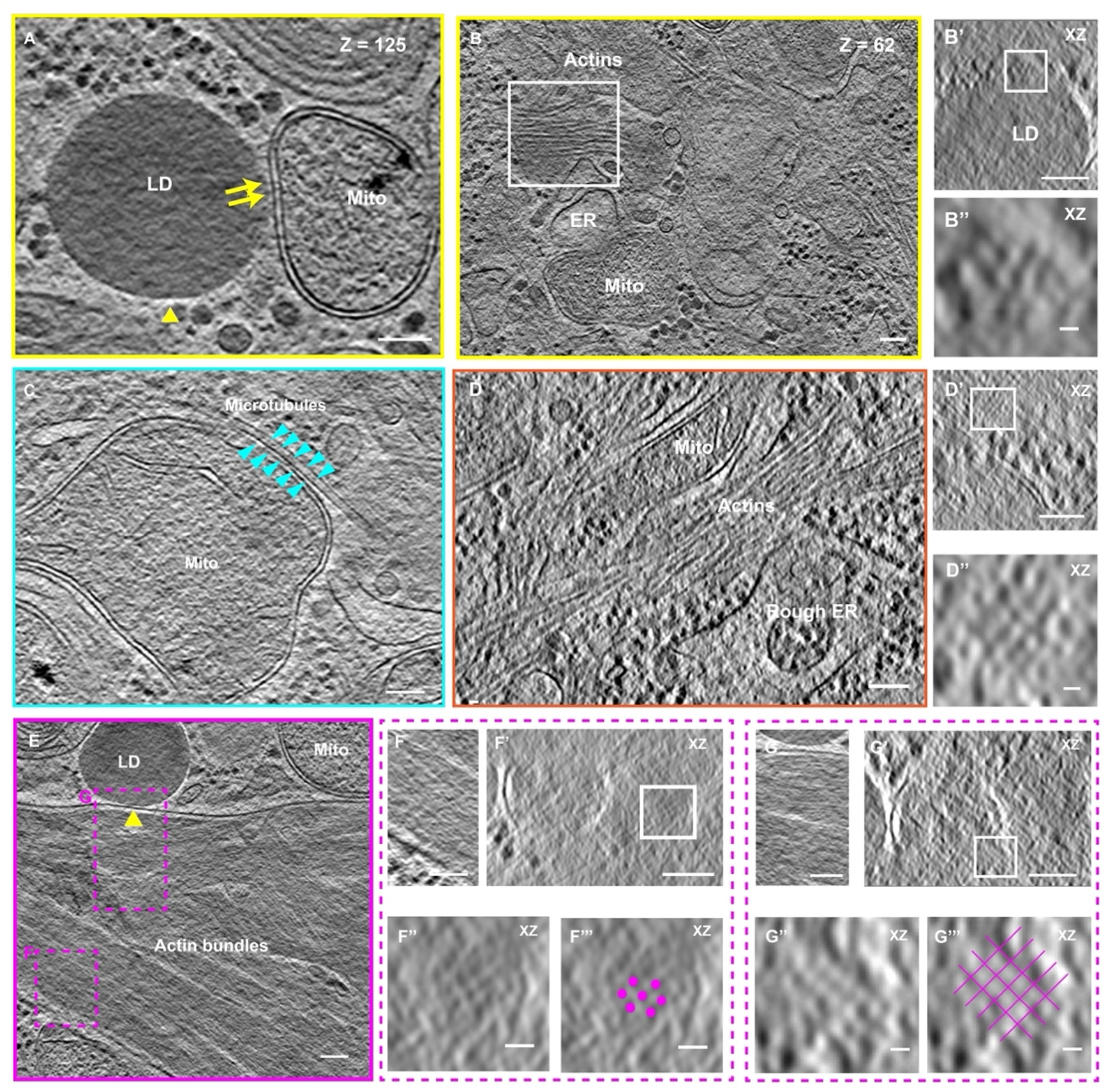
Correlative montage cryo-ET of cytoskeleton arrangements at lipid droplet-mitochondrial contact sites. Tomogram slice and cross-sectional (XZ) views correspond to the color boxed region in Fig. 5H. Yellow boxed region in Fig. 5H notes protein-like densities at the lipid droplet-mitochondrial contact sites (A, Z-slice = 125)), and the actin arrangement (B-B’’) right above (Z-slice = 62) the same ROI lipid droplet (yellow arrowhead, A) bundled by cross-linkers arranged in hexagonal pattern. (C) Microtubule and microtubule-mitochondrial contacts. (D) Organized actin above a mitochondrion (Z-slice = 74). (E-G) Dense actin filament organization near lipid droplet-mitochondrial contact sites. Pink dashed boxed sub-region (F and G) in (E) correspond to subareas of interest where actin was bundled in hexagonal arrays in dense filamentous arrangements. B’, D’, F’, and G’ are cross-section XZ views of B, D, F, G. B’’, D’’, F’’, G’’’ are magnified views of the white boxed region in B’, D’, F’. Scale bars of 100 nm in A, B, B’, C, D, D’, E, F, F’, G, G’, and 10 nm in B’’, D’’, F’’, F’’’, G’, G’’.

**Fig. S6.**
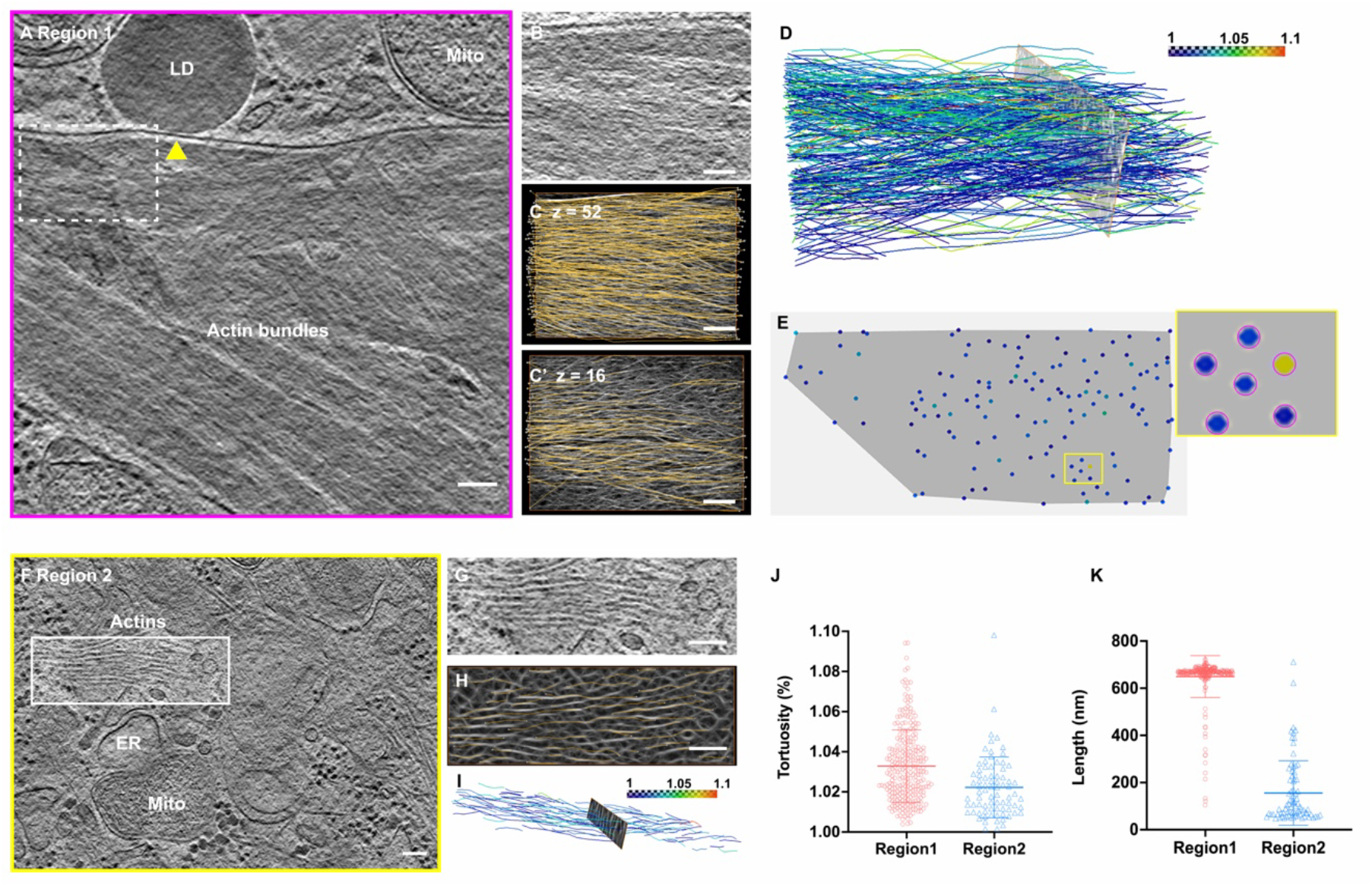
Distinct morphologies of actin-like filaments near the LD-Mitochondrial site captured by MPACT. Actin-like filaments display densely packed (A-E, Region 1) and loosely packed (F-I, Region 2) arrangements. (A) A 20 nm thick tomographic slice through a sub-cellular volume from the LD targeted (yellow arrowhead) FIB-milled lamella collected by 3x3 MPACT (Fig. 4H, pink boxed). (B) An enlarged view of the white boxed region in (A). (C-C’) Overlay of actin tracing and segmentation at different Z-slices from the volume. (D-E) Corresponding segmentation of actin filaments color coded from blue to red based on the tortuosity. (E) One cross-sectional segmentation view show hexagon-like actin packing arrangements observed in Fig. S5 (delineated by pink circles in the zoomed-in view). (F) A 20 nm thick tomographic slice through a sub-cellular volume from the targeted FIB-milled lamella collected by 3x3 MPACT (Fig. 4H, yellow boxed). An enlarged view (G) of the white boxed region in (F) and the corresponding actin segmentation overlay with the tracing (H) and color-coded display based on tortuosity (I) show the loosely packed morphology. (J-K) Statistical characterization of the dense (red) and loose (blue) actin-like filaments around the LD-Mito sites to show the differences in tortuosity (J) and length (K). Scale bars of 100 nm in A-H.

